# Genome-wide association study of cocaine self-administration behavior in Heterogeneous Stock rats

**DOI:** 10.1101/2025.07.17.660811

**Authors:** Montana Kay Lara, Lieselot L. G. Carrette, Thiago Missfeldt Sanches, Oksana Polesskaya, Alicia Avelar, Angela Beeson, Hassiba Beldjoud, Brent Boomhower, Molly Brennan, Denghui Chen, Lindsey China, Apurva S. Chitre, Dana Elizabeth Conlisk, McKenzie Fannon, Benjamin B. Johnson, Elaine Keung, Adam Kimbrough, Jenni Kononoff, Angelica Renee Martinez, Lisa Maturin, Khai-Minh Nguyen, Alex Morgan, Joseph Mosquera, Dyar Othman, Sonja L. Plasil, Jarryd Ramborger, Paul Schweitzer, Sharona Sedighim, Osborne Seshie, Kokila Shankar, Benjamin Sichel, Sierra Simpson, Lauren Cassandra Smith, Elizabeth A. Sneddon, Lani Tieu, Nathan Velarde, Selene Zahedi, Leah C. Solberg Woods, Marsida Kallupi, Giordano de Guglielmo, Abraham A. Palmer, Olivier George

## Abstract

**Background:** Cocaine use disorder (CUD) is a major public health crisis with detrimental individual and societal effects. The specific genes mediating CUD remain largely unknown.

**Methods:** We conducted a genome-wide association study (GWAS) using outbred N/NIH Heterogeneous Stock (HS; n = 836, female = 415, male = 421) rats. We examined multiple CUD-related phenotypes that captured acquisition of self-administration, escalation of intake, and compulsive-like responding.

**Results:** Consistent with the existing literature, these traits were phenotypically and genetically correlated and exhibited modest heritability (h^2^ = 0.07 – 0.16). We identified six genome-wide significant associations. One locus on chromosome 19 was associated with the variable time between cocaine infusions (post infusion interval) and contains several carboxylesterase genes that are orthologous to the human *CES1* gene; notably, carboxylesterases metabolize cocaine. Three non-synonymous coding variants in the genes *Ces1c* and *Ces1d* were in perfect linkage disequilibrium with this locus, suggesting that one or more of them might be the causal SNP. The other 5 loci also contained promising coding and expression variants, including *Trak2,* a gene previously associated with CUD in human GWAS and *Slc10a7*, *Plcl1*, and *Satb2* which have been associated with alcohol and tobacco use disorder.

**Conclusions:** This is the largest genetic study of cocaine self-administration ever conducted in any species. Our results replicate previous loci associated with CUD in humans and provide several novel biological insights including the potential of pharmacological strategies targeting carboxylesterases for the treatment of CUD.

## Introduction

Cocaine use disorder (CUD) is a pressing public health issue. The 2022 National Survey on Drug Use and Health found that approximately 5 million individuals in the US reported using cocaine in the past year ^1^. Notably, 1.4 million people meet the criteria for CUD. The complexity of CUD is further amplified by its frequent comorbidity with other neuropsychiatric disorders, underscoring the challenges in developing effective therapeutic strategies.

CUD is a highly heritable and genetically complex trait. Twin studies estimate the heritability of cocaine dependence to be as high as 70% ^2–4^, a finding supported by recent comprehensive reviews ^5,6^. Complementing these findings, genome-wide association studies (GWAS) have also revealed a significant heritable component, with single nucleotide polymorphism (SNP)-based heritability estimated at 27-30% ^7,8^. Despite the strong evidence of the genetic etiology of CUD, few genome-wide significant associations have been reported: *FAM53B, NCOR2* ^9^, *LINC01411*, *TRAK2*, *LPHN2* ^10^, and *FAM78B* ^11^. The paucity of significant and replicated associations for CUD limits our understanding of this disorder, hampering our ability to identify novel pharmacological targets.

Progress in human studies has been constrained by small sample sizes, complex diagnostic criteria, and the need for cocaine-exposed controls. Rodent models provide a complementary resource for exploring the genetic basis of CUD-like behaviors under tightly controlled experimental conditions. Inbred mouse strains and the Collaborative Cross (CC) recombinant inbred strains exhibit widely varying locomotor response to cocaine^12,13^, and recent studies using the hybrid mouse diversity panel (HMDP) and BXD recombinant inbred mouse population have identified candidate genes associated with cocaine self-administration ^14,15^, including *Fam53b,* an ortholog to the human gene (*FAM53B*) implicated in CUD ^16^.

In addition to inbred strains and recombinant inbred panels, outbred rodent models offer a valuable tool for dissecting the genetic basis of CUD-like behaviors. Outbred populations such as the Heterogeneous Stock (HS) rats provide a high degree of genetic diversity and low levels of linkage disequilibrium, enabling greater mapping precision. Prior work has established the phenotypic diversity of HS rats across a broad range of addiction-relevant behaviors, including cocaine self-administration ^17–24^. Importantly, GWAS conducted in HS rats have identified numerous genetic variants associated with clinically relevant phenotypes related to a broad spectrum of human disorders ^25–32^.

Building on this framework, we expanded the sample used for a phenotypic analysis in de Guglielmo et al. (2024) to include nearly 900 HS rats that had undergone extended access to cocaine intravenous self-administration. This procedure captures key stages of the addiction process, such as initial acquisition of cocaine self-administration, escalation of cocaine intake, motivation to seek cocaine, and compulsive-like cocaine use despite adverse consequences ^33–35^. By dissecting both the macrostructure, such as patterns of intake over multiple sessions, and the microstructure of cocaine self-administration, such as the differences in time between drug infusions, we can examine both correlated and orthogonal aspects of cocaine self-administration. This level of detail is unobtainable in human studies, highlighting a major advantage of performing this work in rats. We then used SNP genotypes at millions of markers to perform a GWAS analysis. In addition to the GWAS results, we used various secondary analysis strategies, such as integration with coding variants and eQTLs to uncover novel genetic drivers of cocaine self-administration behaviors.

## Methods and Materials

To measure behaviors related to CUD, HS rats underwent experimental testing for extended access to cocaine self-administration (Figure 1A). More detailed experimental protocols can be found at https://www.protocols.io/workspaces/george-lab and are also described in prior publications ^22,23^. All phenotypic and genetic data can be found in the supplementals and at UCSD Library Digital Collections (https://doi.org/10.6075/J0QN675X).

**Figure 1.**
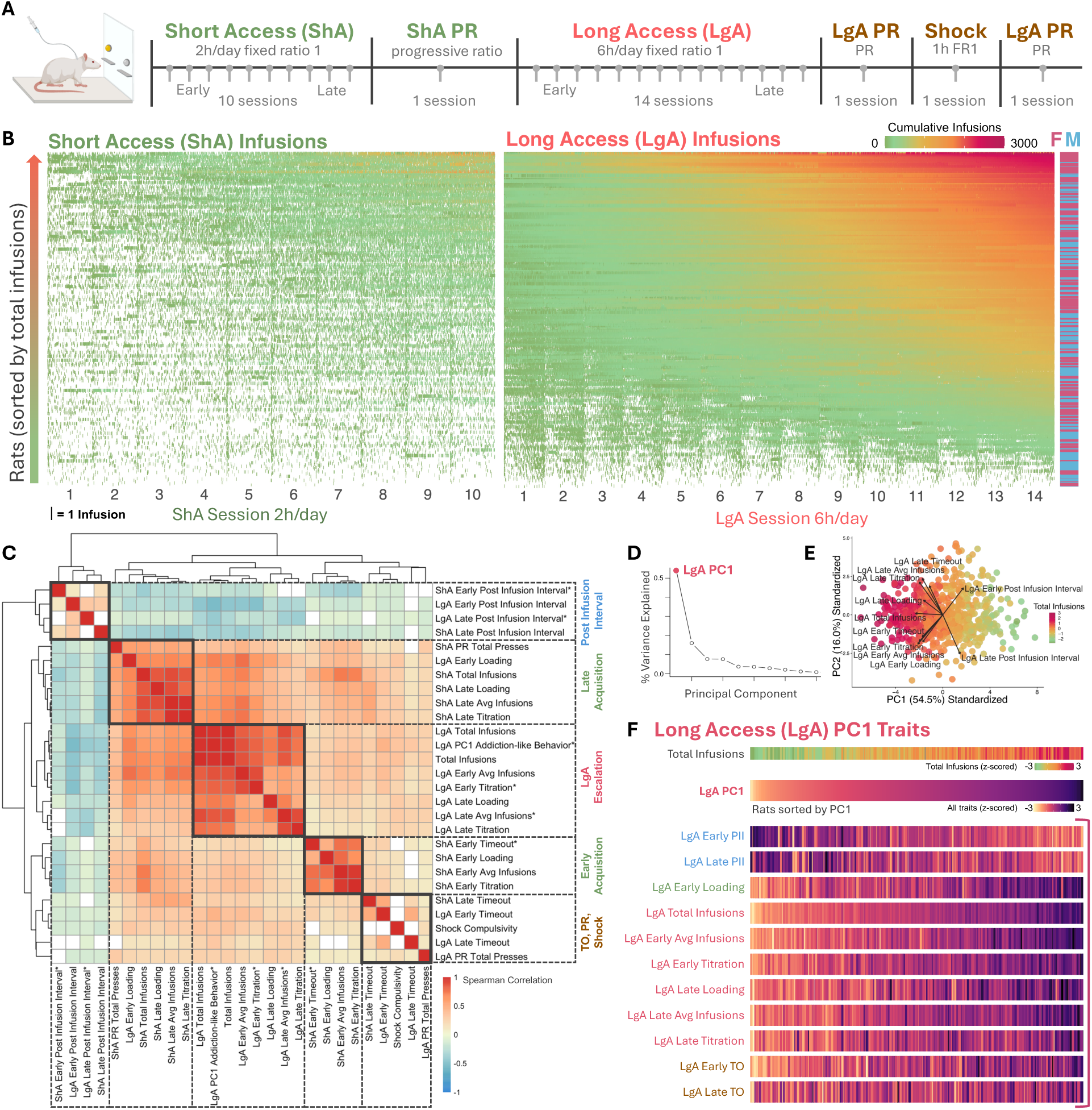
HS rats show individual differences in addiction-like behaviors. A) Timeline of extended access to cocaine self-administration, starting with short access (ShA) for 10 sessions 2 hours per day, then progressive ratio (PR) for 1 session, long access (LgA) for 14 sessions 6 hours per day, PR for 1 hour session, footshock punishment for 1 hour session, and a final PR 1 hour session. ShA and LgA traits were categorized as “early” or “late”, corresponding to the beginning or end sessions of ShA or LgA. B) Individual infusions are shown over time for each rat. The y-axis of all rats is sorted by total infusions. The x-axis shows the sessions for both short and long access. One line indicates one infusion. The color of the infusion line reflects the cumulative infusions the rat has received at that timepoint. C) Behavioral correlations were calculated for all traits using Spearman’s correlation and traits were clustered using average linkage hierarchical clustering. Clusters are indicated with boxes and labeled based on the traits they contain. Cluster label colors are consistent with color labels for the behavioral experiment they came from, with the post infusion interval being notably separated. Non-significant values are indicated in white. Generally, LgA traits clustered together, and both early and late post infusion intervals clustered separately. D) To aggregate traits for a composite behavioral phenotype of addiction-like behavior, we used PC1 of all LgA traits, which accounted for 55% of the variance. E) Biplot of PC1 and PC2 showing where each LgA trait falls along the PC axes. Each dot represents a rat, and the color reflects the total number of infusions. F) All traits included in LgA PC1 addiction-like behavior are represented, and for each heatmap, the x axis is all rats sorted by their LgA PC1 addiction-like behavior value. The total infusions trait is included for reference, and generally, rats with higher total infusions are towards the right of the x axis. Labels for traits included in LgA PC1 addiction-like behavior are also colored by the behavioral cluster in which they belong.

### Animals

This study contains data from 20 cohorts of HS rats (approximately 1,125 total experimental rats), spanning over 8 years. The first 17 cohorts were bred at Wake Forest University School of Medicine by Dr. Leah Solberg Woods (NMcwiWFsm:HS #13673907, RRID:RGD_13673907). The last 3 cohorts were bred at the University of California San Diego by Dr. Abraham A. Palmer (McwiWfsmAap:HS #155269102, RRID:RGD_155269102). Rats were shipped or transferred to Dr. Olivier George’s lab in batches of 30-60 rats at 3 to 4 weeks of age. RFID chips were inserted into all rats at weaning. The first 6 cohorts were tested at the Scripps Research Institute, and the subsequent 14 cohorts were tested at the University of California San Diego. Each cohort consisted of a similar number of female and male rats. Upon arrival at the George lab, all rats from Wake Forest were kept in quarantine for two weeks. Following quarantine or transfer, rats of the same sex were pair-housed in the vivarium with reverse 12-hour light/dark cycle, temperature and humidity control, and ad libitum access to food and water. All experimental testing was approved by the Institutional Animal Care and Use Committees at the Scripps Research Institute or the University of California San Diego. All rats were treated in compliance with the National Institutes of Health Guide for the Care and Use of Laboratory Animals.

### Intravenous catheterization

The detailed protocol for intravenous jugular catheterization can be found at https://www.protocols.io/view/intravenous-jugular-catheterization-for-rats-36wgq5z45gk5/v1. Rats (age 7-9 weeks) were anesthetized with isoflurane vapor (induced at 5% and maintained at 2-3%). The jugular vein was exposed via an incision, and a catheter was inserted through an incision in the back and threaded under the skin to the neck. The catheter was inserted into the vein, secured with sutures, flushed, and checked for blood flow before the surgical site was closed. Post-operative care included administration of antibiotics (Cefazolin 330 mg/kg intramuscular) and analgesics (Flunixin 2.5 mg/kg subcutaneous) administered immediately following surgery and as needed over the next 5 days. Rats were monitored before returning to home cages. After catheterization surgery, rats were allowed to recover for at least one week before testing. Catheters were flushed daily with heparinized saline (10 U/ml of heparin sodium; American Pharmaceutical Partners, Schaumberg, IL, USA) in 0.9% bacteriostatic sodium chloride (Hospira, Lake Forest, IL, USA), which contained 52.4 mg/0.2 ml of Cefazolin).

### Cocaine self-administration (escalation)

The detailed protocol for cocaine self-administration can be found at https://www.protocols.io/view/cocaine-iv-self-administration-dm6gpwx3dlzp/v1 ^22,23^. All behavior testing was performed during the dark cycle, with the start of the session aligned to the lights going off in the housing room. The operant chambers for self-administration experiments were housed in sound-attenuating cubicles (Med Associates, St. Albans, VT, USA). The chambers were equipped with two retractable levers (active and inactive), and sessions began when these were extended. Infusion pumps were used to deliver cocaine HCl dissolved in 0.9% sterile saline at a dose of 0.5 mg/kg per infusion. Rats were weighed each week and given the appropriate corresponding drug concentration for their weight. Responses on the right (active) lever activated the infusion pump, which delivered 0.1 mL of drug over 6 seconds followed by a 20 second timeout. The timeout period was signaled by illumination of a light above the active lever, and rats would not receive additional drug for active lever presses during this time. Responses on the left (inactive) lever were recorded but had no consequence.

After one week of recovery after surgery, rats (age 8-10 weeks) underwent 10 sessions of short access (ShA) to cocaine self-administration for 2 hours per day, 5 days a week, under a fixed ratio 1 schedule (FR1). After ShA, rats underwent 14 sessions of long access (LgA) to cocaine self-administration for 6 hours per day, 5 days a week (Figure 1A).

### Brevital test

After LgA, in order to test catheter patency, rats were administered Brevital (methohexital sodium; 2 mg/kg), which is a barbiturate with short-acting anesthetic effects. Rats with patent intravenous catheters show an immediate loss of muscle tone after Brevital administration. Any rat that failed to react to the Brevital infusion was recorded as a “catheter fail”. If a rat had a catheter fail, behavioral data and health and experimental notes of the rat were reviewed by the team to determine the last good experimental session day where the catheter was functional. If this was unclear, the rat was excluded from subsequent trait analysis. These decisions were made by highly trained individuals who were blind to the genotype of the rat in question.

### Progressive ratio (motivation)

At the end of both ShA and LgA, rats underwent one and two sessions, respectively, of self-administration on a progressive ratio (PR) schedule of reinforcement (Figure 1A). During PR, response requirements for receiving a single dose of cocaine increased in the following manner: 1, 2, 4, 6, 9, 12, 15, 20, 25, 32, 40, 50, 62, 77, 95, 118, 145, 178…^36^. The PR measure of total presses was defined as the number of active lever presses a rat made (not the number of infusions they received). The session ended once a rat did not complete the required number of presses within a 60 minute period.

### Foot shock (compulsive-like responding)

Between the two sessions of PR after LgA, rats underwent a 1-hour foot shock punishment test under the same FR schedule as ShA and LgA self-administration (Figure 1A). Rats received a foot shock (0.3 mA, 0.5 s) for 30% of cocaine infusions over this hour. The “shock compulsivity” measure was calculated as the total number of rewards in this session after the initial shock. Rats who never received a shock were not included in this trait for downstream genetic analysis.

### Phenotype data processing

Behavioral data, including time stamps for active and inactive lever presses and infusions, was recorded by MED-PC IV software. Experimental notes, including any catheter disconnections, sickness, or death, were also recorded. Of the 1,125 rats that underwent behavioral testing, 158 (13%) were excluded due to sickness, death, catheter failure, or other experimental complications, and 131 (11%) were excluded due to missing or unusable genotype data. Thus, data for part or all of the study was available from 836 rats.

Data from MED-PC files were extracted using custom Python scripts. Data were visually inspected by at least 3 people by plotting all summary data for each day for each individual rat. Missing data from catheter disconnections or computer issues was imputed using Soft-Impute ^37^. Specifically, we only imputed values for the total number of infusions for short or long access days if the data was missing. We did not impute timestamp data (list of time signatures for infusions or lever presses), or any other behavioral experiment. Imputation used the number of infusions for behavioral data, but did not consider any genetic data. If a rat died during the experiment, data for the days after death were considered missing and were not imputed or used in subsequent behavioral or genetic analysis. In cases of death or sickness, the team visually inspected all behavioral data to determine the likely onset of sickness or cause of death. Only sessions in which the animal appeared healthy were included in the trait analysis. To avoid including data potentially influenced by early signs of illness or experimental complications leading to death, any sessions during the four days immediately preceding death were always excluded, as well as any subsequent sessions after the onset of sickness. All data that passed quality checks, including imputed data, was used in the genetic analysis.

If only a portion of data was kept for a rat, the missing data was denoted with NAs. While we were able to impute total number of infusions on a given day of short or long access, missing timestamp data could not be imputed, thus some calculated traits that rely on timestamp data could not be determined. After applying these quality control measures data from 836 rats were available for the subsequent genetic analysis, however due to missing data that could not be imputed, for some traits, fewer than 836 rats were available for analysis.

### Addiction-like trait calculations

All traits are detailed in the supplementals. For both ShA and LgA, early and late denote if a trait was derived from the first 3 or the last 3 or 4 sessions, respectively. Average infusions is the mean number of infusions over the designated sessions. Escalation refers to the increase in the total number of infusions over LgA, and specifically LgA late average infusions is the measure that is most often associated with this trait. Post infusion interval is the median time in between infusions over the designated sessions. Loading is the total number of infusions received in the first 10 minutes over the designated sessions. Titration is the total number of infusions received in the last hour over the designated sessions. Timeout is the average difference between the active lever presses and number of infusions over the designated sessions. PR total presses refers to the total number of active lever presses made in the designated session. For calculated traits that take the sum of infusions across multiple sessions, rats with missing data for any of the days within the time period specified were excluded (e.g., if a rat had NA for session 14 because it was sick, LgA late traits was not calculated). For any trait that was calculated using timestamps (e.g., traits that require total or average numbers of lever presses or infusions), rats without timestamp data were excluded. For shock measures, rats that never received a foot shock (even if they received an infusion) were excluded. The total number of infusions were calculated for both ShA and LgA, and these were summed to produce a grand total infusion measure.

### Behavioral correlations and PCA

To examine the correlations between all trait measures, phenotypic correlations were determined using Spearman’s correlation. These correlations were visualized using Seaborn Clustermap, and hierarchical clustering was used to group traits according to their patterns of similarity. We performed principal component analysis (PCA) on all calculated traits from LgA (11 measures in total), including early and late post infusion interval, average infusions, loading, titration, and timeout, as well as LgA total infusions. These traits spanned multiple behavioral clusters. The first principal component (PC1) explained 54% of the variance of these traits; we termed this PC “LgA PC1 addiction-like behavior” and included it in downstream genetic analysis.

### Trait processing

To ensure reproducibility, all traits used for GWAS were processed in the same way. First, we used linear regression to remove the effects of any covariate that explained more than 2% of the variance, including age, cohort, sex, coat color, or weight. Then all traits were quantile normalized as part of our pre-processing GWAS pipeline (https://github.com/sanchestm/GWAS-pipeline). This process addresses any deviations from normality in the traits and allows for the use of the same GWAS significance thresholds for all traits.

### Genotyping

To genotype HS rats, we used low-coverage whole genome sequencing. DNA was obtained from spleens, which were collected postmortem, as described in detail by Chen et al. ^38^. Detailed protocols for spleen collection, DNA isolation, sample processing, preparing sequencing libraries, and performing the sequencing can be found on protocols.io here: https://www.protocols.io/workspaces/cgord/publications. DNA was extracted using the DNA Advance kit and sequencing libraries were prepared using the Twist 96-Plex Library Prep kit (TWIST Bioscience). Reads were aligned to the rat reference genome mRatBN7.2 from the Rat Genome Sequencing Consortium (GCA_015227675.2 GCF_015227675.2) using BWA-mem v0.7.17. Mapped sequences were used to construct haplotypes and impute biallelic single nucleotide polymorphism (SNP) genotypes using STITCH v1.6.6 ^39^. This process resulted in a high-quality set of 7,358,643 SNPs with imputation quality scores (INFO) ≥ 0.9. We additionally removed all SNPs with high missing rates (missing rate > 0.1; 23,389 removed), low minor allele frequencies (MAF 0 - 0.005; 1,887,082 removed, most of which were monomorphic), and extreme deviations from Hardy-Weinberg Equilibrium (HWE p < 1e-10; 2,604 removed). This filtered set of 5,446,333 SNPs in 836 HS rats was used for all downstream analyses.

### Genetic analysis

We performed a genome-wide association study on the traits described in the prior section using the mixed linear model analysis (MLMA) function from the Genome-wide Complex Trait Analysis (GCTA) software package ^40^ as described previously ^26,29,31^. Briefly, the GWAS pipeline (https://github.com/sanchestm/GWAS-pipeline) uses a genetic relatedness matrix (GRM), which estimates the genetic relationship between individuals based on their genotypes on all autosomes, to control for population structure and accurately determine the genetic contribution to the measured traits. We used the GCTA leave one chromosome out (LOCO) method to avoid proximal contamination ^41^. SNP heritability estimates were calculated with GCTA using the GRM and a restricted maximum likelihood (REML) approach ^40^. GWAS significance thresholds were calculated using 1,000 permutation tests ^42^, which we were able to use across all traits because all traits were quantile normalized, making the genotype and phenotype distributions identical. SNPs reached genome-wide significance according to the following threshold: –log_10_(*p*) = 5.58 (α = 0.05).

## Results

### HS rats exhibit wide variation in cocaine self-administration

Traits were derived from the following experimental assays: short access (ShA) and long access (LgA) to cocaine self-administration, progressive ratio (PR), and shock (Figure1A). Rats exhibited substantial individual differences in cocaine intake both during ShA and LgA (Figure 1B). While some rats maintained low levels of intake (sorted at the bottom of both ShA and LgA intake graphs in Figure 1B), others exhibited dramatic escalation of intake, some exceeding over 150 infusions per session by the end of LgA. This pattern shows the development of pronounced differences in cocaine intake under both short and long access conditions, which broadly parallels the variability in cocaine use among humans.

To characterize distinct behavioral domains underlying the range of addiction-like traits measured, we performed correlation and cluster analysis (Figure 1C). Almost every pairwise comparison in the matrix was significant. This pervasive covariance replicates our previous results ^22^ and suggests that the animals’ intake patterns, pauses, effort, and shock tolerance reflects a common behavioral construct of addiction-like behavior and are likely at least partially controlled by a common set of heritable and environmental factors. Despite this pervasive covariance, the dendrogram still resolved into several coherent sub-modules. Emergent behavioral clusters with highly correlated traits included early and late acquisition (with mostly ShA-derived measures) and LgA escalation (with mostly LgA-derived measures). The post infusion interval cluster, which reflects the time between infusions, included both ShA and LgA post infusion interval measures, suggesting that it may be a stable trait with a high genetic component. This cluster was moderately anti-correlated with other behavioral clusters. This relationship is expected because a rat with a high intake will spend less time between infusions. Finally, timeout (TO), PR, and shock were clustered together, suggesting that they may represent similar measures of motivation to seek cocaine despite adverse consequences, including lack of access to cocaine (TO), increased effort (PR), and punishment (foot shock).

### Principal component analysis of cocaine self-administration

To capture a composite measure of addiction-like behavior, we performed principal component analysis (PCA) on all traits derived from LgA sessions because these traits exhibited the greatest variability. The analyzed traits spanned multiple behavioral domains from the cluster analysis, including early and late measures of average intake, escalation of intake, and post infusion intervals. The first principal component (PC1) explained 55% of the variance among these traits (Figure 1D), providing a robust aggregate measure of addiction-like behavior. The biplot of PC1 and PC2 (Figure 1E) shows the direction of each trait along these axes, and the continuum of z-scored total infusions along PC1. Additionally, the heatmaps of all LgA traits (Figure 1F) highlight the relationship between individual LgA traits and LgA PC1 addiction-like behavior. Furthermore, traits from different behavioral domains were well-represented (indicated by the behavioral cluster coloring), demonstrating that PC1 captures diverse aspects of LgA cocaine intake.

### GWAS

To identify genetic loci associated with cocaine self-administration behaviors, we performed GWAS on all measures from ShA, LgA, PR, and shock (Supplemental Report and Supplemental Table 1). Of these 27 traits, there were six genome-wide significant associations (−log_10_(*p*) = 5.58, α = 0.05) between traits and regions across the HS rat genome. The porcupine plot in Figure 2A summarizes the combined p-values for all 6 traits with significant associations, including ShA early post infusion interval, LgA late average infusions, ShA early timeout, LgA late post infusion interval, LgA PC1 addiction-like behavior, and LgA early titration. Traits from multiple behavioral clusters contributed to significant associations, indicating that genetic influences span multiple aspects of cocaine self-administration.

**Figure 2.**
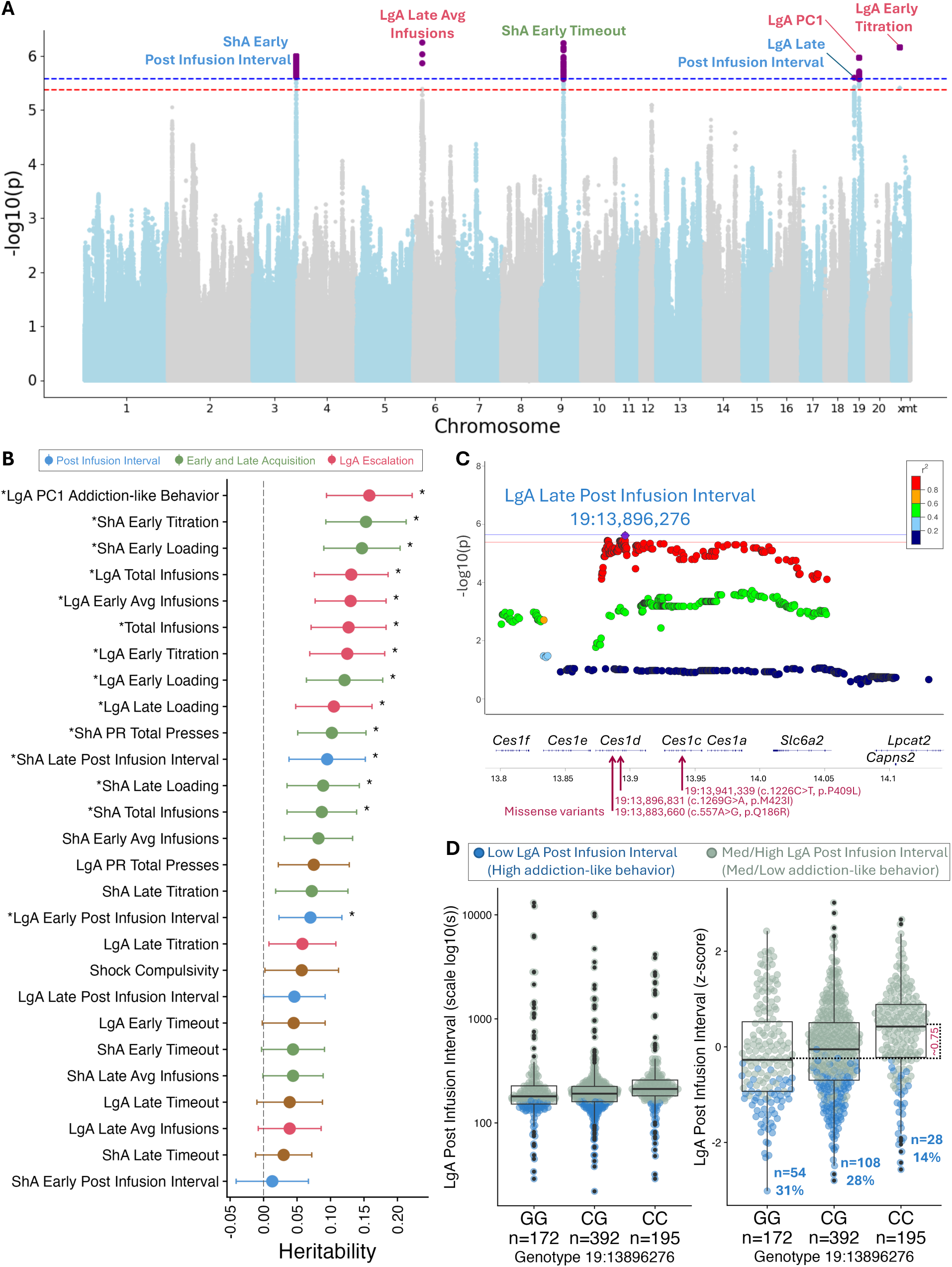
Genome-wide association study for cocaine self-administration in HS rats. A) The porcupine plot displays all *p*-values combined for traits with a significant association distributed across the entire genome of HS rats. The orange and yellow lines show the significance thresholds derived from the permutation tests: −log_10_(*p*) = 5.58 (α = 0.05) and −log_10_(*p*) = 5.38 (α = 0.10), respectively. B) Heritability of all cocaine self-administration traits. Error bars represent standard error; the color reflects the behavioral cluster from which the trait belongs (Figure1B). Stars indicate significant heritabilities (p < 0.05). C) Regional association plot zooming into the locus on chromosome 19 that was significantly associated with post infusion interval. The individual dots represent SNPs, and the color of the dots indicate linkage disequilibrium between the top SNP (in purple) and other nearby SNPs. The blue and red lines show the significance thresholds: −log_10_(*p*) = 5.58 (α = 0.05) and −log_10_(*p*) = 5.38 (α = 0.10), respectively. Below the x-axis shows the genes mapped to that chromosomal position. There are three missense variants located in this locus. D) Effect plot for Post Infusion Interval depicting the genetic effect of the top SNP, with genotype on the x-axis, including homozygous alternate alleles, heterozygous, and homozygous reference alleles. Both plots show LgA post infusion interval on the y-axis, with the first being log_10_ scaled, and the second z-scored for better visualization. The z-scored difference (∼0.75) of Post Infusion Interval between CC and GG genotypes is indicated between the dashed lines. The blue dots represent rats who were in the bottom quartile for LgA Post Infusion Interval.

### Heritability

To quantify the proportion of phenotypic variance attributable to SNPs, we calculated SNP-based heritability for each trait. SNP heritability estimates reflect the extent to which variation in the trait is explained by additive effects of all the SNPs in the HS rat genome. Heritabilities for all traits ranged from low to moderate (Figure 2B, Supplemental Table 2). The highest significant heritability (h^2^ = 0.16, *p* = 0.005) was observed for LgA PC1 addiction-like behavior. Additionally, multiple traits across behavioral domains for ShA and LgA showed significant heritability ranging from h^2^ = 0.07 to 0.16 (*p* < 0.05). For example, the total intake of cocaine infusions during LgA was heritable (LgA total infusions, h^2^ = 0.13, *p* = 0.009), which is a measure that is often used across drug self-administration studies to determine resistance or vulnerability to the development of addiction-like phenotypes.

### Locus for LgA late post infusion interval

GWAS identified a significant locus on chromosome 19 associated with LgA late post infusion interval, a measure reflecting the time elapsed between successive cocaine infusions. The top SNP within this locus was located at position 19:13,896,276 (-log_10_(p) = 5.61), and the associated linkage disequilibrium (LD) block spanned from ∼13,800,000 bp to ∼14,050,000 bp, a region that is approximately 0.25 mb (Figure 2C). Three non-synonymous coding variants were identified within this locus that were in almost perfect LD (r^2^ > 0.99) with the top SNP. Two of these were missense variants located in the coding region of the *Ces1d* gene: 19:13,883,660 (c.557A>G, p.Q186R) and 19:13,896,831 (c.1269G>A, p.M423I). The third missense variant, 19:13,941,339 (c.1226C>T, p.P409L), was in the coding region of the *Ces1c* gene. The genes *Ces1c* and *Ces1d* encode carboxylesterase 1 enzymes and are orthologous to human *CES1*. The enzyme Carboxylesterase 1 is one of the main enzymes responsible for the metabolism of cocaine ^43^. Based on the available data, it is not clear whether these coding variants would be expected to increase or decrease metabolism of cocaine.

In addition to coding variants, several expression quantitative trait loci (eQTL) were identified within this LD block.^20^ These eQTLs affect expression of *Rasd2* in the nucleus accumbens core, prelimbic cortex, and whole brain tissue (all with r^2^ = 1.0). *Rasd2* has been implicated in several relevant phenotypes, including sensitivity to the stimulant effects of cocaine ^44^. Finally, the coding region for the gene *Slc6a2,* which encodes the norepinephrine transporter, which is inhibited by cocaine, is also located within this locus; however, we did not identify any coding variants or eQTLs for this gene, suggesting that it is not the cause of this association.

The top SNP in this locus had a minor allele frequency of 48%. Rats with the CC genotype at this SNP exhibited lower LgA post infusion intervals (Figure 2D). Lower post infusion interval values, or those that are in the bottom quartile, correlated with high levels of cocaine intake and greater addiction-like traits.

### Loci for additional cocaine self-administration traits

GWAS identified five additional loci exceeding the genome-wide significance threshold that were associated with CUD-like traits (Figure 2A, Supplemental Report). All significant loci, their corresponding genes in range, eQTLs, and splice QTLs (sQTLs) are listed in Supplemental Tables 3-6, respectively.

LgA PC1 addiction-like behavior, derived from LgA traits spanning multiple behavioral domain clusters, was significantly associated with a locus on chromosome 19 at 29,567,757 bp (-log_10_(p) = 5.98). Two eQTLs (r^2^ = 0.9) were identified in whole brain tissue and the nucleus accumbens core for the gene *Lsm6*, which is involved in RNA binding and splicing. Additionally, four sQTLs for *Lsm6* were identified in the basolateral amygdala, nucleus accumbens core, prelimbic cortex, and whole brain tissue (r^2^ = 0.65, 0.94, 0.66, and 0.68, respectively). Finally, the gene *Slc10a7*, which has been implicated in smoking ^10,45,46^ and nicotine dependence ^47^ was located in this locus.

ShA early post infusion interval, which represents the median time elapsed between infusions during the first three sessions of ShA, was associated with a locus on chromosome 3 (Supplemental Report). The top SNP within this locus was at position 3:163,216,140 (-log_10_(p) = 5.997) (Supplemental Report). Seven missense variants in perfect LD (r^2^ = 1.0) with this SNP were identified within the gene *Zfp831,* which encodes a zinc finger protein. This region also included an eQTL for *Zfp831* in the basolateral amygdala. Additionally, three eQTLs for the gene *Gnas,* which encodes the stimulatory alpha subunit of a G protein, were detected at SNPs in moderate to high LD (r^2^ = 0.6 – 1.0) in the lateral habenula, nucleus accumbens core, and the prelimbic cortex. This locus also contained a coding variant for the gene *Ctsz* (r^2^ = 1.0), as well as eQTLs for *Ctsz* in the nucleus accumbens core and whole brain tissue (both with r^2^ = 1.0).

LgA late average infusions, a measure of escalation of intake, was associated with a locus on chromosome 6, with the peak SNP at position 6:34,284,564 (-log_10_(p) = 6.25) (Supplemental Report). Six significant coding variants in perfect LD (r^2^ = 1.0) were identified in this locus, all in the gene *Rad51ap2*, which plays a role in *Rad51* activity as part of DNA repair pathways and meiotic recombination ^48^. Two additional SNPs in high LD with the top SNP (r^2^ = 0.73 and 0.75) were sQTLs for the gene *Vsnl1* in brain and prelimbic cortex tissues. *Vsnl1* encodes a protein that is part of the visinin/recoverin subfamily of neuronal calcium sensor proteins ^49^.

Lastly, ShA early timeout, calculated as the difference between the number of active lever presses and the number of infusions received, captures burst pressing and possibly perseverative behavior. A locus on chromosome 9 was significantly associated with ShA early timeout, with the peak SNP at position 9:57520529, and the implicated region spanned approximately 4 mb (Supplemental Report). While there were no coding variants or eQTLs in this region that were in high LD (r^2^ > 0.7) with the top SNP, several SNPs in this locus were in moderate LD (0.5 < r^2^ < 0.7) with the top SNP and are associated with genes that have been implicated in human GWAS for alcohol drinking, nicotine, and cocaine use: *Plcl1, Satb2,* and *Trak2* ^10,45,46^.

## Discussion

To investigate the underlying genetic factors influencing resilience or susceptibility to CUD-related traits, we performed a GWAS in HS rats using an extended access self-administration model. The phenotypes assessed in this study capture key dimensions of addiction-like behaviors. The extended access model allowed us to characterize escalating intake, increased motivation to take the drug, and compulsive-like behavior despite negative consequences ^22,34^. We identified multiple significant loci associated with CUD-like behaviors, notably including a region containing coding variants for *Ces1c* and *Ces1d* that encode enzymes involved in cocaine metabolism that is associated with the post infusion interval.

Twin studies of cocaine use, abuse, and dependence have high narrow-sense heritability estimates ranging from ∼0.3 – 0.7 ^2–4,50,51^. More recent GWAS studies estimate SNP-based heritability for cocaine dependence to be around ∼0.3 ^7,8^. SNP heritability represents the proportion of phenotypic variance explained by the additive effects of SNPs included in the study. SNP heritability is expected to be lower than heritability estimates from twin studies in part because it only captures variance attributable to common variants. We found heritabilities for CUD-like traits in HS rats ranged from 0.07 to 0.16. Interestingly, the highest heritability was attributed to the principal component derived from multiple CUD-like behavioral domains: LgA PC1 addiction-like behavior. This is notable because PC1 integrates multiple phenotypic dimensions into a robust metric, and the relative high heritability of 16% underscores the utility in mapping using principal components. These findings suggest that genetic influences on addiction-like behavior may become more pronounced as drug exposure escalates, aligning with the progressive nature of substance use disorders in humans.

Human GWAS of CUD remain in their infancy, with sample sizes still numbering in the thousands. These studies are limited by the difficulty of obtaining cases as well as uncertainty about whether controls should have previous exposure to cocaine. Genome-wide significant associations have been reported for a few genes including *FAM53B, NCOR2* ^9^; *FAM78B* ^11^; and *LINC01411*, *TRAK2*, and *LPHN2* ^10^. One of the genes we identified (*Trak2*) is the rat homolog of *TRAK2*, which is one of the few genes that has also been associated with CUD in humans ^10^. The fact that some of the genes are not observed across species likely reflects low power in both human and rat GWAS, differences between the CUD phenotype in humans and the phenotypes measured in HS rats, and differences in common, polymorphic alleles present in the human populations that have been studied and HS rats. Given these limitations, the fact that even a single gene has been found in both species is remarkable.

In addition to the overlap of *Trak2* between rats and human CUD GWAS, several other genes we identified have also been identified in human GWAS of other SUDs. For example, *SLC10A7* has also been implicated in smoking initiation ^46^ and nicotine dependence ^47^; *PLCL1* has been implicated in the maximum number of alcoholic drinks (Kapoor et al., 2013) and smoking initiation ^45,46^; and *SATB2* has been implicated smoking initiation ^46^. We identified a genome-wide significant locus on chromosome 19 for LgA post infusion interval, which contained several *Ces1* genes. Three missense variants were present in *Ces1c* and *Ces1d*, which encode enzymes that are involved in the metabolism of various substances, including cocaine ^43,52^. As a drug-metabolizing enzyme, CES1 hydrolyzes cocaine into inactive metabolites, influencing both the duration and intensity of cocaine’s effects on the brain. At present, it is not clear whether or how these coding variants might alter enzymatic activity. Differences in *Ces1* activity in HS rats may impact CUD-like traits, reflecting the role of this gene in regulating the response to repeated drug exposure. This locus was significantly associated with the elapsed time between infusions, a key behavioral measure linked to the compulsive nature of cocaine self-administration, as hypothesized by the compulsion zone theory ^53^, which proposes that cocaine self-administration is controlled by pharmacokinetic constraints, where rats maintain cocaine levels above the priming/remission threshold but below the satiety threshold to sustain drug-taking behavior. *CES1* has already been discussed as a potential target for therapeutic interventions ^54^, and our findings provide further support for studies aimed at modulating addiction-like behaviors by modulating CES1 activity.

In addition to drug metabolism, we identified genes of various biological function that were associated with addiction related traits, as well as expression and splice QTLs that may explain how these loci affect the behavioral differences seen across rats ^20^. Central to the process of addiction are neurocircuits involved in reward, motivation, learning, and inhibitory control ^55–58^. Within the 6 significant loci we identified in this study, several genes had eQTLs and sQTLs in brain regions heavily involved in these processes ^20^. Genes in the implicated loci with significant eQTLs and sQTLs were located in the nucleus accumbens (*Rasd2, Gnas, Ctsz, Lsm6*), which is involved in the dopamine reward circuit associated with binging and intoxication, as well as the prelimbic cortex (*Rasd2, Gnas, Vsnl1, Lsm6*) and amygdala (*Zfp831, Lsm6)*, which play a role in negative emotional states and stress responses in the withdrawal stage ^55,56,59–64^. Critically, these circuits involve dopamine signaling, which can be altered as a result of chronic drug use ^65–71^. Among the genes identified in this study, several may be involved in this process. For example, *Rasd2,* which encodes a Ras protein enriched in the striatum, has been shown to modulate dopamine neurotransmission, and have effects on motor response after cocaine exposure ^44,72,73^. *Gnas*, which encodes the stimulatory alpha subunit of G protein coupled receptors, including dopamine receptors, has been shown to regulate dopamine receptor signaling and have downstream effects after cocaine exposure ^74–77^. Lastly, *Trak2,* which encodes the trafficking kinesin protein 2, interacts and transports GABA receptors in the brain ^78,79^. The GABAergic system is crucial for inhibitory neuronal activity that regulates dopamine reward and reinforcement pathways in addiction ^80^.

There is growing evidence that there are shared genetic drivers underlying multiple substance use disorders (SUDs) or addiction-related behaviors ^81,82^. Much of this genetic signal appears to represent non-substance specific factors that may reflect behavioral under-control ^83^, which may partially act through their ability to influence initiation of drug taking. In contrast, the procedure used in our study assured that all rats have an opportunity to try cocaine. Therefore, our GWAS is intended to capture substance-specific genetic factors. The identification of *Ces1c* and *Ces1d*, which influence cocaine metabolism, reflect this bias towards cocaine-specific factors.

Many prior animal studies have examined responses to drug exposure. For examine, Zhou, et al. ^84^, examined gene expression at the single cell level using some of the same HS rats that are included in this GWAS. That study highlighted differences in gene expression that were observed after 4 weeks of cocaine abstinence in rats that self-administered either large or small amounts of cocaine. In contrast, the present study sought to identify genes that predispose rats to voluntarily self-administer large or small amounts of cocaine. These two approaches highlight the difference between temporally variable states (cocaine exposure) and stable traits (genotype).

To our knowledge, this is the largest study of extended access to cocaine self-administration in outbred rats. Our GWAS identified multiple loci significantly associated with CUD-like traits in HS rats, including escalation of intake of cocaine, time elapsed between cocaine infusions, and an aggregate measure of addiction-like behavior. This study highlights the utility of rich behavioral phenotyping and using genetically diverse rats to identify novel genetic loci associated with the multifaceted process of drug use and the transition to addiction-like behaviors. Samples from some of the rats used in these studies are available thought the Cocaine biobank (https://www.cocainebiobank.org/) ^23^. Genes associated with CUD in humans remain limited, however our GWAS identified one gene (*Trak2*) that has also been identified by human GWAS of CUD, and the novel identification of *Ces1* offers a fresh avenue for future studies.

## Acknowledgements

Funding was provided by the following NIH grants: P50DA037844, P30DA060810, U01DA051234, and U01DA043799.

## Author contributions (CRediT)

Conceptualization and funding: AAP and OG. Writing original draft: MKL. Writing reviewing and editing: MKL, LLGC, OP, BBJ, LCSW, AAP, and OG. Investigation: LLGC, AA, HB, BB, MB, LC, DEC, MF, AK, JK, ARM, LM, KN, AM, JM, DO, SLP, JR, SS, KS, BS, SS, LCS, EAS, LT, NV, SZ, MK, GD. Data curation: MKL, LLGC, TMS, OP, MB, EK, AM, DO, BS, AAP, and OG. Methodology: TMS, DC, ASC, BBJ. Project administration: LLGC, OP, and PS. Supervision: LLGC, OP, MK, GD, AAP and OG. Resources: AB, OS, and LCSW. Analysis: MKL, LLGC, TMS, OP, and EK.

